# m^6^A mRNA Methylation Regulates Early Pancreatic β-Cell Differentiation

**DOI:** 10.1101/2023.08.03.551675

**Authors:** Sevim Kahraman, Dario F De Jesus, Jiangbo Wei, Natalie K. Brown, Zhongyu Zou, Jiang Hu, Chuan He, Rohit N Kulkarni

## Abstract

N^6^-methyladenosine (m^6^A) is the most abundant chemical modification in mRNA, and plays important roles in human and mouse embryonic stem cell pluripotency, maintenance, and differentiation. We have recently reported, for the first time, the role of m^6^A in the postnatal control of β-cell function in physiological states and in Type 1 and 2 Diabetes. However, the precise mechanisms by which m^6^A acts to regulate the development of human and mouse β-cells are unexplored. Here, we show that the m^6^A landscape is dynamic during human pancreas development, and that METTL14, one of the m^6^A writer complex proteins, is essential for the early differentiation of both human and mouse β-cells.

## INTRODUCTION

N6-methyladenosine (m^6^A) is reported to be the most prevalent post-transcriptional modification in mRNA and non-coding RNA and widespread across various tissues (Liu et al., 2020). m^6^A mRNA modification is installed co-transcriptionally by the methyltransferase complex consisting of at least METTL3, METTL14, and WTAP proteins and removed by the eraser proteins such as FTO and ALKBH5 (Fu et al., 2014). FTO has been shown to have m^6^A-independent roles in gene regulation (Kim et al., 2022), while data from human malignancies have demonstrated that ALKBH5 has contradictory functions, behaving as an oncogene in some cancers and a tumor suppressor in others (Qu et al., 2022). The YTH family of reader proteins recognize and bind the m^6^A-modified RNAs and mediate post-transcriptional modification by affecting multiple stages of mRNA metabolism, such as nuclear export, alternative splicing, mRNA stability, or translation (Wang et al., 2022).

The differential m^6^A methylation of transcripts has been observed in various human diseases (He and He, 2021) including diabetes (De Jesus et al., 2019, 2023). We have previously reported decreased levels of m^6^A modifications in pancreatic islets of established Type 1 (De Jesus et al., 2023) and Type 2 Diabetes patients (De Jesus et al., 2019), and identified a number of hypomethylated genes that are critical for β-cell biology and associated with the development of diabetes.

m^6^A methylome analyses performed on major fetal human tissues revealed differential methylation among different tissue types, and highlighted the tissue-specific developmental role of this mRNA modification (He et al., 2023; Wei et al., 2022; Xiao et al., 2019). Likewise, several studies have reported the involvement of m^6^A-dependent mRNA regulation in cellular development processes such as hematopoiesis (Vu et al., 2017), neurogenesis (Yoon et al., 2017), adipogenesis (Kobayashi et al., 2018), or spermatogenesis (Lin et al., 2017). However, the contributions of m^6^A RNA modifications to the genesis of pancreatic endocrine cells and the consequent impact on maintenance of glucose homeostasis in the body are not fully explored.

We have reported that deletion of Mettl14 in postnatal pancreatic β-cells leads to a decreased number of β-cells and leads to development of diabetes in mice (De Jesus et al., 2019). Others have confirmed these observations and shown that depletion of Mettl3 and Mettl14 specifically in pancreatic β-cells caused hyperglycemia in 14-day old mice (Wang et al., 2020). These studies argue that m^6^A mRNA modifications are important for β-cell biology and function. However, the mechanistic role of m^6^A modifications in regulating the early development of human β-cells remains largely unclear, in part because of the difficulty in accessing high quality human fetal β-cells. Improvements in the protocols for in vitro differentiation that mimic human pancreas development coupled with the availability of new tools has enabled investigators to study how m^6^A modifications contribute to the process of β-cell differentiation.

In the present study, we analyzed human fetal β-cells and β-like cells derived from human pluripotent stem cells and observed that m^6^A modulators are expressed in pancreatic β-cells undergoing development. To understand the landscape and dynamics of m^6^A modification in β-cell development, we performed RNA sequencing (RNA-seq) and m^6^A MeRIP-sequencing (m^6^A-seq) at different stages of in vitro β-cell differentiation. We profiled the changes in m^6^A modifications and expression levels of key transcription factors important for β-cells to evaluate their characteristics. Finally, we employed three different Cre-recombinase mouse models to determine the impact of Mettl14 ablation during different stages of pancreatic development.

Our findings reveal the dynamic nature of the m^6^A landscape throughout human pancreas development, highlighting the crucial role of METTL14, a component of the m^6^A writer complex, during early differentiation of β-cells in both humans and mice.

## RESULTS

### METTL14 Levels Increase in Developing Human Pancreatic **β**-cells

We began by evaluating the expression of m^6^A modulators during islet cell maturation. All m^6^A regulators were highly expressed in human pancreatic α-cells and β-cells and several were altered in mature compared to fetal β-cells (Figure 1A). These changes appeared to be more dynamic in β-cells (Figure 1A) compared to α-cells (Figure 1B). Among the m^6^A writer complex proteins, *METTL14* expression showed a significant increase in adult compared to fetal β-cells (Figure 1A). Immunostaining of human adult and fetal pancreas sections (patient information in Supplementary Table 1) revealed highly abundant levels of METTL14 protein in late fetal (28-41 w) and adult β-cells (0-65 y) compared to early fetal β-cells (12-18 w) (Figure 1C, D) suggesting enhanced expression of the writer protein during development. Since in vitro differentiation is a useful tool to study stages of in vivo human islet cell development, we re-analyzed datasets from embryonic stem cell derived β-cells (Veres et al., 2019) and again observed altered gene expression levels of m^6^A regulators during in vitro β-cell differentiation (Figure 1E). Consistently, *METTL14* expression was particularly increased in more advanced stages of β-cell differentiation (e.g. stage 6 (S6) β-like cells), compared to the premature stage (e.g. S5 endocrine progenitors; Figure 1E). We also confirmed that METTL14 protein levels gradually increased during the in vitro differentiation of H1 and MEL1, two human embryonic stem cell lines (hESCs), into β-like cells (Figure 1F, H). Consistent with the increase in expression levels of the m^6^A “writer” gene *METTL14* during β-cell differentiation, global m^6^A levels were increased with differentiation of both H1 and MEL1 hESCs towards S6 β-like cells, supporting a role for m^6^A mRNA modification during development (Figure 1I). Transcriptome and immunostaining analysis of human fetal β-cells together with in vitro β-like cell differentiation studies showed dynamic changes in METTL14 levels and again supported the involvement of m^6^A modification during β-cell development.

**Figure 1.**
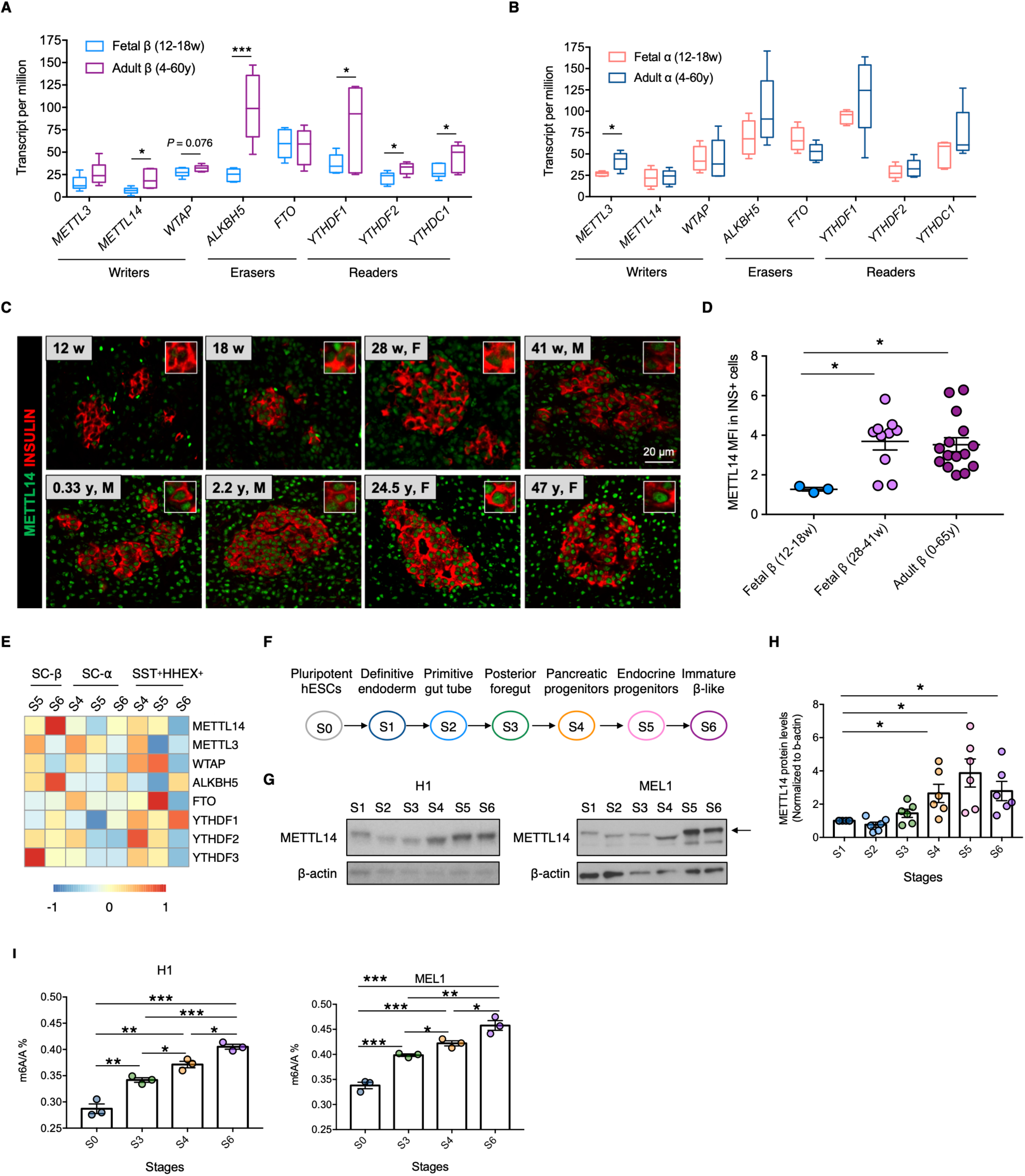
Expression of m^6^A modulators is dynamic during development of the human pancreas. **(A)** Re-analysis of RNA-seq data (Blodgett et al., 2015) showing expression levels of genes involved in m^6^A RNA methylation in fetal β-cells (12-18 weeks, n = 6 independent biological samples) and adult β-cells (4-60 years, n = 6 independent biological samples). Two tailed t-test. **(B)** Expression levels of genes in fetal α-cells (12-18 weeks, n = 5 independent biological samples) and adult α-cells (4-60 years, n = 5 independent biological samples). Two tailed t-test. **(C)** Representative images of immunostaining of human pancreatic sections collected from cadaveric donors. Top panel (gestational ages between 12 weeks to 41 weeks) and bottom panel (0-65 years). METTL14 (green) and insulin (red). **(D)** Mean fluorescence intensity (MFI) of METTL14 staining in insulin+ area in human pancreas sections. Early fetal β (12-18 weeks, n = 3 independent biological samples), late fetal β (28-41 weeks, n = 10 independent biological samples), adult β (0-65 years, n = 15 independent biological samples). **(E)** Re-analysis of single cell RNA-seq data (Veres et al., 2019) showing expression levels of m^6^A regulators in SC-β, SC-α, and SST+HHEX+ cells. **(F)** Stages of in vitro differentiation of hESCs into β-like cells. **(G)** Representative WB images showing changes in METTL14 protein levels during in vitro differentiation in H1 (left panel) and MEL1 hESC lines (right panel). Arrow shows METTL14 protein. **(H)** Changes in METTL14 protein levels during differentiation. H1 n = 3 and MEL1 n = 3 independent biological samples. Multiple t-test followed by Holm-Sidak. **(I)** Changes in percentage of global m^6^A levels during in vitro differentiation in H1 (left panel) and MEL1 hESC lines (right panel).

### Human Pancreas Development is Characterized by Dynamic m^6^A Landscape Remodeling

We performed RNA-seq and m^6^A-seq at four different stages to profile changes in mRNA modifications during development to specifically study the biological relevance of enhanced m^6^A during in vitro β-cell differentiation. Successful differentiation of hESCs into posterior foregut (S3), pancreatic progenitors (S4), and immature β-like cells (S6) was confirmed by the upregulation of stage-specific marker genes (Figure S1A-C). As expected, m^6^A peaks were mostly enriched in the 3’untranslated region (3’UTR) and near stop codons (Figure S1D-F). The PCA plot showed that samples were clustered according to differentiation stages (Figure 2A).

**Figure 2.**
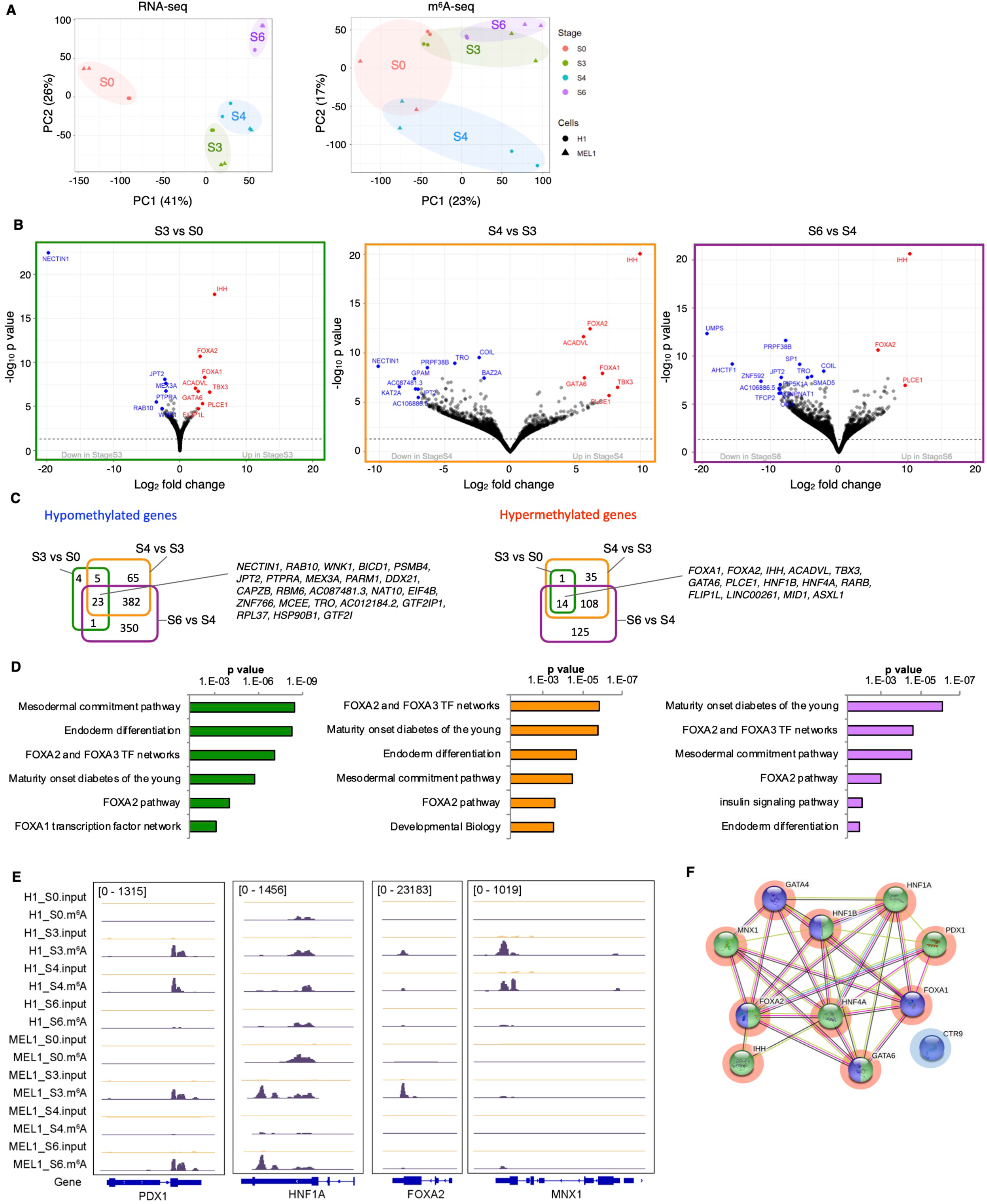
RNA N^6^-methyladenosine sequencing reveals dynamic changes in m^6^A decoration during in vitro **β**-like cell differentiation. **(A)** PCA plot of RNA-seq (left) and m^6^A-seq (right) after regressing out batch effect in H1 and MEL1 hESC line (H1 n = 2 and MEL1 n = 2 independent biological samples). **(B)** Volcano plot showing hypomethylated and hypermethylated genes at S3 vs S0 (left), S4 vs S3 (middle), and S6 vs S4 (right) (H1 n = 2 and MEL1 n = 2 independent biological replicates). **(C)** Venn diagrams showing number of hypomethylated (FC ≤ −2) and hypermethylated (FC ≥ 2) genes (FDR < 0.1) at S3 vs S0, S4 vs S3, and S6 vs S4 (H1 n = 2 and MEL1 n = 2 independent biological samples). **(D)** Pathway analyses of hypermethylated genes at S3 vs S0 (left), S4 vs S3 (middle), and S6 vs S4 (right) (H1 n = 2 and MEL1 n = 2 independent biological samples). **(E)** Coverage plots of m^6^A peaks in the PDX1, HNF1A, FOXA2, MNX1 genes showing H1 and MEL1 S0, S3, S4, S6 (H1 n = 2 and MEL1 n = 2 independent biological samples). **(F)** Protein interaction analysis of hypermethylated genes involved in pancreas development (green) and endoderm differentiation (purple). Red circled genes are upregulated (FDR < 0.1, FC ≥ 2), and blue circled genes are downregulated (FDR < 0.1, FC ≤ −2) at S3 vs S0, S4 vs S3, and S6 vs S4.

While RNA-seq clearly segregated the stages, there was considerable variability in early stages of β-cell differentiation followed by a more defined m^6^A methylome at later stages. Interestingly, S4 samples clustered at a distance from S3 and S6 samples, indicating the unique methylome that reflects stage S4 may represent a transition between stages.

We identified 59 differently methylated sites in 51 genes by comparing stage S3 vs S0, as well as 739 differently methylated sites in 626 genes comparing S4 vs S3, and 1205 differently methylated sites in 981 genes comparing S6 vs S4 [false discovery rate (FDR) < 0.1] (Figure 2B). *NECTIN1, JPT2, RAB10, TRO* are among the most significantly hypomethylated transcripts [FDR < 0.1, fold change (FC) ≤ −2], and *FOXA1, FOXA2, IHH, ACADVL, TBX3, GATA6, PLCE1* are among the most significantly hypermethylated transcripts (FDR < 0.1, FC ≥ 2) that are common at each stage of differentiation (Figure 2C). Pathway analyses of the transcripts affected by m^6^A hypermethylation revealed several networks that are important for β-cell development such as “endoderm differentiation,” “maturity-onset diabetes of the young,” and “FOXA2 and FOXA3 transcription factor network,” comparing S3 vs S0, S4 vs S3, or S6 vs S4 (Figure 2D, Supplementary Dataset 1). We then intersected m^6^A-seq with RNA-seq to understand how differential m^6^A peaks influence the transcriptomics of cells during in vitro differentiation, since it has been reported that extensive transcriptional control is necessary to activate or suppress switch genes that influence cell fate during directed β-cell differentiation (Weng et al., 2020). The intersection revealed that the hypermethylated transcripts which are involved in endoderm differentiation and pancreas development such as *HNF1A, HNF1B, HNF4A, FOXA1, FOXA2, GATA4, GATA6, PDX1, and MNX1* (Figure 2E) were also upregulated at S3 vs S0, S4 vs S3, and S6 vs S4 (FDR < 0.1, FC ≥ 2) (Figure 2F, and Supplementary Dataset 2). These data suggest that m^6^A modifications contribute to regulating the expression of these key genes involved in β-cell development.

### *METTL14* Controls Endoderm Differentiation in Mice and Humans

In our previous study, we reported that deletion of Mettl14 in postnatal pancreatic β-cells leads to changes in the β-cell identity and numbers of β-cells and leads to the development of diabetes after birth in mice (De Jesus et al., 2019). Given the phenotypic perturbations observed in β-cells of the KO mice, we hypothesized that m^6^A mRNA modifications play a role in β-cell biology and development. To address this hypothesis, we first successfully depleted Mettl14 in mouse induced pluripotent stem cells (miPSCs) using siRNA (Figure 3A). Spontaneous differentiation of Mettl14 knockdown miPSCs into embryoid bodies (EBs) (Figure 3A) showed striking impairment in the generation of endoderm lineage compared to scramble treated mouse EBs (Figure 3B). We next explored the requirement of METTL14-mediated m^6^A RNA methylome in hESC differentiation by simultaneously inducing METTL14 knockdown in H1 or MEL1 hESCs with doxycycline (Dox) treatment and differentiating the hESCs towards the three embryonic germ layers using EB formation assays (Figure 3C). It has been shown that depletion of m^6^A methyltransferase complex in mouse and human ESCs affects expression levels of the pluripotency gene Nanog and impairs ESC exit from self-renewal (Batista et al., 2014). In order not to disturb the pluripotency capacity of hPSCs before initiating the differentiation towards three germ layers, we generated inducible METTL14 knockdown hESC clones and depleted METTL14 during spontaneous differentiation. RNA-seq analysis of Dox-induced METTL14 knockdown EBs (METTL14 iKD1/2/3 Dox+) and control EBs (iSCR Dox+) showed that depletion of METTL14 resulted in the upregulation of 540 genes (FDR < 0.1, FC > 1.5) and downregulation of 344 genes (FDR < 0.1, FC < −1.5) (Figure 3D). Pathway analysis revealed that the upregulated genes are associated with mesoderm and ectoderm lineages including “osteoblast differentiation,” “neural crest differentiation,” and “mesodermal commitment pathway” (Figure 3E and Supplementary Dataset 3). On the other hand, downregulated genes are associated with cell division such as “mitotic G1 phase and G1/S transition” and “nucleosome assembly” (Figure 3E and Supplementary Dataset 3). Increased expression of genes associated with ectoderm and mesoderm lineages – with no significant alterations in expression of genes involved in endoderm lineage upon depletion of METTL14 in EBs (Figure 3F) – suggested that METTL14 deficiency disrupts the balance of hESC differentiation and skews differentiation in favor of ectoderm and mesoderm. One of the factors that affects pluripotency and lineage choice in hESCs is disruptive insulin receptor-mediated signaling (Teo et al., 2021). Considering that METTL3 or METTL14 depletion in cells causes hypomethylation of the transcripts involved in insulin/IGF1-AKT pathway and impairment of insulin signaling (De Jesus et al., 2019), it can be argued that defective insulin signaling could interfere with endoderm differentiation in METTL14-depleted EBs.

**Figure 3:**
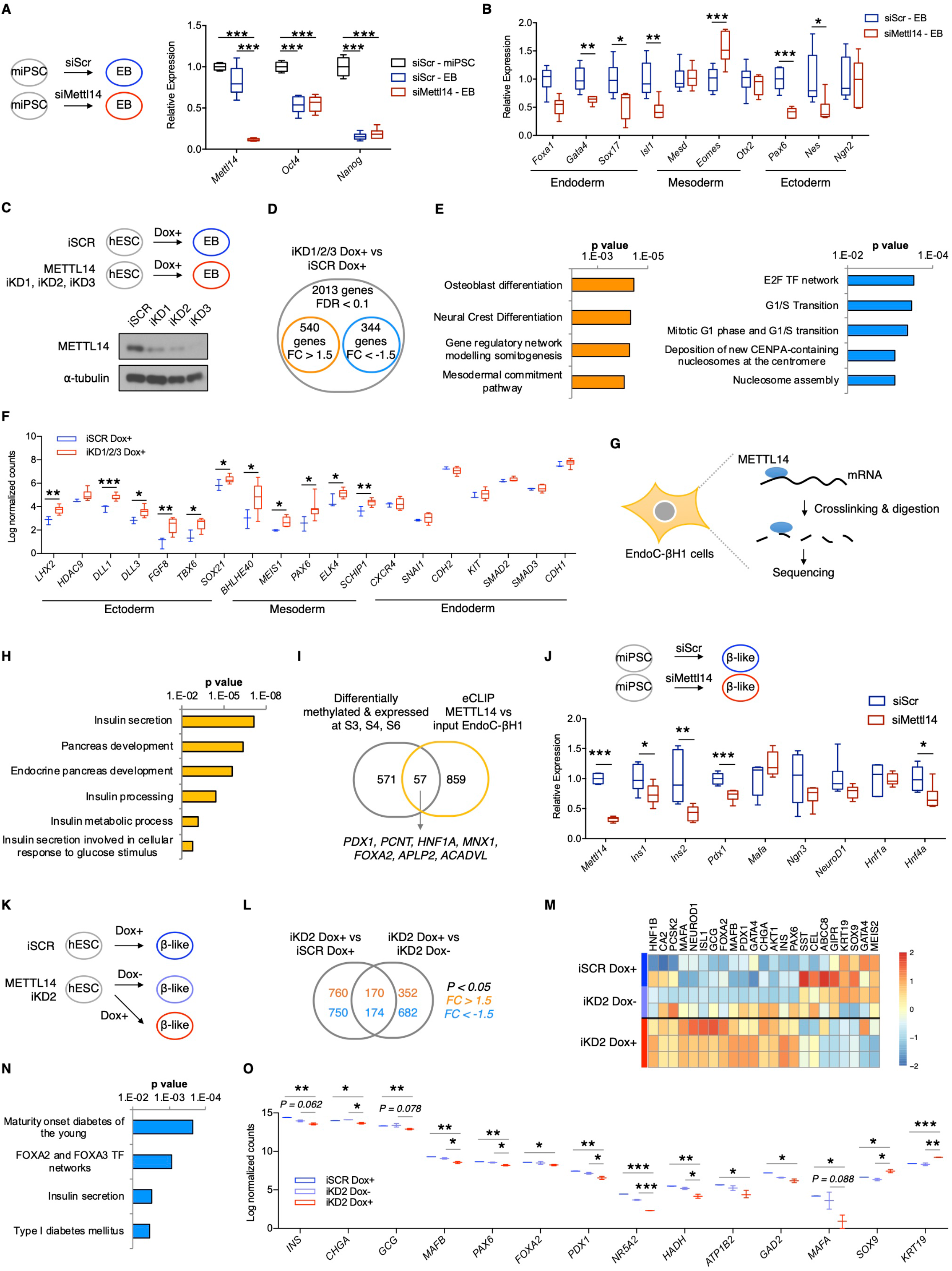
METTL14 controls endoderm differentiation in mice and humans. **(A)** Mouse iPSCs were transiently transfected with siRNAs targeting Mettl14 or scramble as a control and then spontaneously differentiated to embryoid bodies for 4 days. RT-PCR analysis performed on miPSCs (n = 4 independent biological samples) and EBs (siScr EB n = 6 and siMettl14 EB n = 6 independent biological samples). Expression was normalized against the mean level in undifferentiated miPSCs. **(B)** Gene expression levels of markers of endoderm, mesoderm, and ectoderm. Expression was normalized against the mean level in siScramble EBs. siScr EB n = 6 vs siMettl14 EB n = 6 independent biological samples. p values were calculated by two tailed t-test. **(C)** Inducible knockdown of METTL14 was generated using lentiviral vectors in H1 and MEL1 human ESCs. Inducible hESCs were spontaneously differentiated into embryo bodies (EBs) for 10 days and treated with doxycycline to induce shRNA expression. **(D)** RNA-seq of EBs revealed 2013 DEGs (FDR < 0.1) in METTL14 iKD EB (H1 n = 6 and MEL1 n = 3 independent biological samples) vs iSCR EB (H1 n = 2 and MEL1 n = 1 independent biological samples). **(E)** Upregulated (left panel) and downregulated (right panel) pathways in METTL14 iKD EB (H1 n = 6 and MEL1 n = 3 independent biological samples) vs iSCR EB are represented (H1 n = 2 and MEL1 n = 1 independent biological samples). **(F)** Gene expression levels in METTL14 iKD EB cells (H1 n = 6 and MEL1 n = 3 independent biological samples) compared to iSCR EB (H1 n = 2 and MEL1 n = 1 independent biological samples). Multiple t-test. **(G)** eCLIP performed using antibodies specific to METT14 to detect mRNAs bound by METTL14 in EndoC-βH1 β-cells. **(H)** Enriched GO terms for transcripts bound by METTL14 protein. Input n = 2, METTL14 IP n = 2 independent biological samples. **(I)** Venn diagram showing the number of transcripts differentially methylated and expressed at S3, S4, S6 and the number of transcripts bound by METTL14 protein. **(J)** miPSCs were transiently transfected with siRNAs targeting Mettl14 or scramble as a control and then differentiated to β-like cells. Gene expression levels were measured on day 4 when Mettl14 was silenced efficiently. Expression was normalized against the mean level in scramble. siScr n = 6 vs siMettl14 n = 6 independent biological samples. p values were calculated by two tailed t-test. **(K)** H1 hESCs were differentiated to β-like cells in the presence or absence of Dox and insulin-expressing β-like cells were sorted by flowcytometry. **(L)** Venn diagram shows number of genes that are altered by METTL14 knockdown in H1 hESC-derived β-like cells (p < 0.05, FC > 1.5 or FC < −1.5). iSCR Dox+ n = 2, iKD2 Dox-n = 2, iKD2 Dox+ n = 3 independent biological samples. **(M)** Heatmap showing expression levels of genes altered by METTL14 knockdown in H1 hESC-derived β-like cells (iSCR Dox+ n = 2, iKD2 Dox-n = 2, iKD2 Dox+ n = 3 independent biological samples). **(N)** Pathway analysis of downregulated genes in METTL14 iKD2 Dox+ cells compared to iSCR Dox+ H1 hESC-derived β-like cells. iKD2 Dox+ n = 3 vs iSCR Dox+ n = 2 independent biological samples (p < 0.05, FC < −1.5). **(O)** Gene expression levels in METTL14 iKD2 Dox+ cells compared to iSCR Dox+ and iKD2 Dox-H1 hESC-derived β-like cells. iSCR Dox+ n = 2, iKD2 Dox-n = 2, iKD2 Dox+ n = 3 independent biological samples. Multiple t-test.

### METTL14 Controls Early **β**-cell Differentiation

Considering EndoC-βH1 cells are derived from the human fetal pancreatic bud, have not developed fully into their final stages, and express many β-cell markers, we performed enhanced crosslinking and immunoprecipitation (eCLIP) assays to identify m^6^A sites regulated by METTL14 in human fetal β-cells (Figure 3G). We identified 1969 sites in 916 transcripts that are bound by the METTL14 protein. Gene Ontology (GO) analysis revealed that METTL14 protein targets transcripts that are involved in “pancreas development” and “insulin secretion” in β-cells (Figure 4H, Figure S2A). For example, METTL14 binds to several transcripts (*PDX1*, *HNF1A,* and *PCNT*) which are hypermethylated and upregulated at stage S6 vs S4, and downregulated in METTL14 knockdown β-like cells (Figure 3I, Figure S2B-F) suggesting that METTL14-mediated m^6^A modifications play a role in regulation of expression of these genes involved in pancreas and β-cell development.

**Figure 4:**
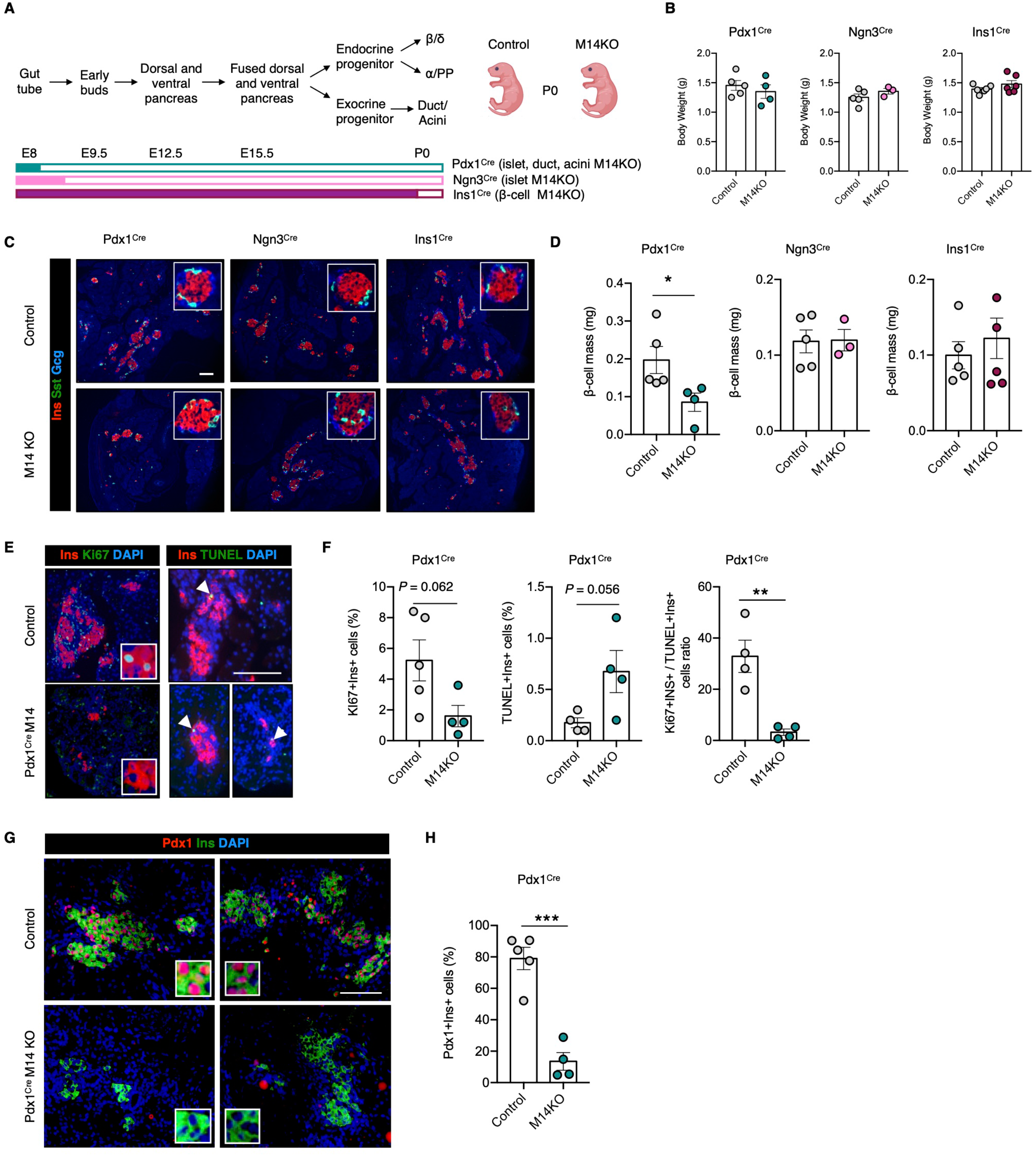
METTL14 controls early **β**-cell differentiation. **(A)** Developmental stages of mouse pancreas and estimated time points for Pdx1, Ngn3, and Insulin promotor-driven Mettl14 depletion during pancreas development. **(B)** Body weight of pups at the time of birth (P0). Pdx1^Cre^ Control n = 5, Pdx1^Cre^ M14 KO n = 4, Ngn3^Cre^ Control n = 5, Ngn3^Cre^ M14 KO n = 3, Ins1^Cre^ Control n = 5, Ins1^Cre^ M14 KO n = 6 independent biological samples. **(C)** Cocktail staining of pancreatic sections (P0). Insulin (red), somatostatin (green), glucagon (blue). Scale bar is 50 μm. Insets are magnified 4 times. **(D)** β-cell mass (P0). Pdx1^Cre^ Control n = 5, Pdx1^Cre^ M14 KO n = 4, Ngn3^Cre^ Control n = 5, Ngn3^Cre^ M14 KO n = 3, Ins1^Cre^ Control n = 5, Ins1^Cre^ M14 KO n = 6 independent biological samples. **(E)** Ins/Ki67 staining (left panel) and insulin/TUNEL staining of pancreatic sections at P0 (right panel). Insulin (red), Ki67 or TUNEL (green), DAPI (blue). Scale bar is 50 μm. Insets are magnified 4 times. Arrowheads show TUNEL+Ins+ cells. **(F)** Quantification of percentage of Ki67+Ins+ cells, TUNEL+Ins+ cells, and ratio of Ki67+Ins+/TUNEL+Ins+ cells in Pdx1^Cre^ P0 pancreas. Pdx1^Cre^ Control n = 5, Pdx1^Cre^ M14 KO n = 4 independent biological samples. **(G)** Pdx1/Ins staining of pancreatic sections at P0. Pdx1 (red), insulin (green), DAPI (blue). Scale bar is 50 μm. Insets are magnified 4 times. **(H)** Quantification of percentage of Pdx1+Ins+ cells. Pdx1^Cre^ Control n = 5, Pdx1^Cre^ M14 KO n = 4 independent biological samples. p values were calculated by unpaired two tailed t-test.

Considering that METTL14 binds to several genes involved in pancreas development (Supplementary Dataset 4) which are also dynamically m^6^A methylated and expressed during in vitro differentiation (Supplementary Dataset 1), we investigated whether depletion of METTL14 perturbs β-cell differentiation. For this purpose, we employed a differentiation protocol to generate immature β-like cells using mouse iPSCs (Figure S3A, B). Mettl14 knockdown during the differentiation of mouse β-like cells resulted in downregulation of *Ins1, Ins2, Pdx1,* and *Hnf4a* genes which argues that Mettl14 is essential for normal β-cell development in mice (Figure 3J). We next asked whether METTL14 is required for human β-like cell differentiation by simultaneously inducing knockdown of the writer protein with doxycycline (Dox) treatment during the differentiation of H1 or MEL1 hESCs towards β-like cells (Figure 3K). Insulin-expressing β-like cells were sorted by flow cytometry and transcriptomic changes in METTL14 knockdown β-like cells (iKD2 Dox+) versus control β-like cells (iSCR Dox+ or iKD2 Dox-) were analyzed by RNA-seq. Depletion of METTL14 during β-cell differentiation resulted in downregulation of several genes that confer β-cell identity such as *INS*, *CHGA*, *PDX1*, *PAX6,* and upregulation of exocrine marker genes such as *KR19* and *SOX9* (Figure 3L, M and Supplementary Dataset 4).

Pathway analysis of genes downregulated by METTL14 knockdown (p < 0.05, FC < −1.5) revealed key networks that are important for β-cell development and function such as “maturity onset diabetes of the young,” “FOXA2 and FOXA3 transcription factor networks,” “insulin secretion,” and “type 1 diabetes mellitus” (Figure 3N, O and Supplementary Dataset 4).

### Mettl14 is Indispensable for In Vivo Pancreatic **β**-cell Development

To validate our findings in an in vivo mammalian system, we next created three different Mettl14 knockout mouse models (M14KO) by crossing Mettl14^fl/fl^ mice with 1) Pdx1^Cre^ mice to knockout Mettl14 in early pancreatic progenitors giving rise to pancreatic islet, ductal, and acinar cells (Hingorani et al., 2003); 2) Ngn3^Cre^ mice to deplete Mettl14 in endocrine progenitors giving rise to pancreatic islet cells (Schonhoff et al., 2004); and 3) Ins1^Cre^ to deplete Mettl14 specifically in pancreatic β-cells (Thorens et al., 2015) (Figure 4A). All M14KO mice, irrespective of their genotype, were born with normal body weight compared to their control littermates (Figure 4B). While Ngn3^Cre^ and Ins1^Cre^ M14KO newborns have comparable β-cell mass, Pdx1^Cre^ M14KO newborns had significantly lower β-cell mass compared to their control littermates (Figure 4C, D). Further investigation of Pdx1^Cre^ M14KO pancreas revealed a decrease in the numbers of proliferating β-cells (Ki67+Ins+) and an increase in apoptotic β-cells (TUNEL+Ins+) which resulted in a significantly reduced ratio of proliferating to apoptotic β-cells in M14KO newborn pancreases (Figure 4E, F). These data suggested that loss of Mettl14 in Pdx1+ pancreatic progenitors during early stages of pancreas development (before or at ∼E8.0) severely impairs the formation of β-cells. Further studies on P0 pancreas sections obtained from Pdx1^Cre^ M14KO mice showed that numbers of PDX1+ β-cells in M14KO newborn pancreases are significantly decreased compared to control littermates (Fig. 4G, H). Together, these data indicate that a lack of the Mettl14 writer protein during early stages of pancreas development (before or at ∼E8.0) affected growth by decreasing numbers of proliferating versus apoptotic cells, leading to a significant loss of Pdx1+ and eventually decreasing β-cell mass in the newborns.

## DISCUSSION

Development of pancreatic β-cells is known to be regulated by the spatial and temporal expression of transcription factors (Conrad et al., 2014; Jennings et al., 2013; Wilson et al., 2003). We have recently reported that METTL14 is required for the maintenance of a functional β-cell mass (De Jesus et al., 2019). In this study, we specifically explored the role of m^6^A in the regulation of both mouse and human β-cell development.

Previous studies examining human fetal pancreas showed that islet-like structures appear by the end of the first trimester (12-13 w) and distinct cell clusters form by week 18 (Jennings et al., 2013; Jeon et al., 2009). Pancreatic islets develop mostly during the second trimester (14-26 w) and remodeling of the developing islets occurs throughout late gestation and after birth (Fowden and Hill, 2001). We report that human fetal β-cells express m^6^A modulators during the first or second trimester of pregnancy and the expression levels of some m^6^A modulators increase after birth. These data imply that m^6^A-mediated post-transcriptional modifications occur during β-cell development. Specifically, dynamic changes in gene and protein expression levels of METTL14 shown by transcriptomics and immunostaining respectively points to the involvement of this writer in β-cell development. Interestingly, the changes in expression of m^6^A modulators during pancreas development are dynamic – as indicated by transcriptomic studies performed on human fetal pancreatic cells showing upregulation of several m^6^A modulators with age in β-cells – compared to there being virtually no significant change in m^6^A modulators in α-cells, indicating a cell specific role of m^6^A-dependent modifications.

Dynamic studies on β-cell development are hampered largely by the lack of availability of human fetal pancreases for research. The availability of improved protocols for stepwise differentiation of pluripotent stem cells into pancreatic β-like cells enabled us to directly study the function of m^6^A modifications during β-cell differentiation (Kahraman et al., 2016). Our methylome and transcriptome analyses demonstrated hypermethylation and upregulation of a number of key transcription factors such as FOXA2, HNF1A, HNF1B, HNF4A, GATA4, GATA6, PDX1, and MNX1 during differentiation. While FOXA2 has been shown to control mRNA levels of PDX1, HNF1A, and HNF4A, mutations in HNF1A, HNF1B, HNF4A, and PDX1 are known to cause maturity onset diabetes of the young (MODY) (Burgos et al., 2021). Mutations in GATA6 are associated with human pancreas agenesis, and mutations in GATA4, and MNX1 cause permanent neonatal diabetes mellitus (Burgos et al., 2021; Teo et al., 2013). Pancreatic defects including reduced number of β-cells or pancreatic agenesis have been reported in humans with mutations in HNF1B, GATA6, PDX1, and MNX1 (Allen et al., 2012; Flanagan et al., 2014; Stoffers et al., 1997; Teo et al., 2016). Likewise, Hnf1b, Gata4/Gata6, Pdx1 or Mnx1-deficient mice fail to develop pancreas of normal size and rather develop diabetes (Flanagan et al., 2014; Haumaitre et al., 2005; Stoffers et al., 1997; Xuan et al., 2012). Expression levels of these pancreatic transcription factors could define the number of β-cells generated during development and therefore also contribute to an individual’s susceptibility to develop diabetes later in life. It is also possible that a small β-mass underlies the poor ability of β-cells to compensate when individuals develop insulin resistance during aging or pregnancy (Meier et al., 2008). Hypermethylation and upregulation of these critical transcription factors at differentiation stages 3, 4, and 6 indicate a temporal involvement of m^6^A in pancreatic endocrine cell genesis.

Previous studies have shown that m^6^A sites act as regulators of definitive endoderm specification of hESCs (Cheng et al., 2022) and that loss of METTL14 during differentiation of stem cells affects the lineage choice (Batista et al., 2014). Our in vitro depletion studies in both mouse and human cells demonstrate that the deficiency of METTL14 impairs development of β-cells. In line with the in vitro findings, we provide in vivo evidence that m^6^A modifications are important for β-cell development and that Mettl14 is also required for proper development of the mouse pancreas. While depletion of Mettl14 in endocrine progenitors (Ngn3^Cre^ M14KO) or in pancreatic β-cells (Ins1^Cre^ M14KO) does not affect β-cell mass after birth, depletion of Mettl14 in early pancreatic progenitors (Pdx1^Cre^ M14KO) leads to a reduced β-cell mass already at P0. Thus, Pdx1^Cre^ M14KO newborns start their life with a compromised β-cell mass due to fewer proliferating cells and an increase in the number of apoptotic cells. Together, these data imply that the temporal regulation of mRNA methylation by Mettl14 is crucial in determining β-cell mass in mammals.

m^6^A modifications have been known to regulate gene expression levels by affecting multiple stages of mRNA metabolism, such as nuclear export, alternative splicing, mRNA stability, or translation (Wang et al., 2022). Therefore, depletion of Mettl14 could impair post-transcriptional regulation of transcripts that are dynamically methylated and expressed during pancreas development. For example, it has been shown that MafA mRNA stability was decreased by depletion of Mettl3/Mettl14 writer complex in MIN6 mouse β-cells and failure to maintain MafA mRNA stability impacted functional maturation of neonatal murine β-cells (Wang et al., 2020). Another study showed that m^6^A decorations regulated mRNA decay of SOX2, which is an important gene for the definitive endoderm specification of hESCs (Cheng et al., 2022). Although additional studies are necessary to explore the underlying mechanisms of loss of Pdx1+ β-cells leading to decreased β-cell mass, the present study sheds lights into the dynamic changes in m^6^A decorations on key transcripts during human β-cell differentiation and provides insights into the significance of METTL14 during β-cell development both in vitro and in vivo.

## ACKNOWLEDGEMENTS

We thank the Network for Pancreatic Organ donors with Diabetes (nPOD; RRID:SCR_014641), a collaborative type 1 diabetes research project supported by JDRF (nPOD: 5-SRA-2018-557-Q-R) and The Leona M. & Harry B. Helmsley Charitable Trust (Grant#2018PG-T1D053, G-2108-04793) for providing human fetal and adult pancreatic sections. We thank Dana-Farber/Harvard Cancer Center supported in part by an NCI Cancer Center Support Grant # NIH 5 P30 CA06516 for the use of the Specialized Histopathology Core, which provided human fetal pancreatic sections. We thank Dr. Andrew B. Leiter for sharing his Tg(Neurog3-cre)C1Able mouse. Flow cytometry experiments were performed in the Joslin Flow Cytometry Core, supported by the Diabetes Research Center (DRC) (nos. P30DK036836 and S10 OD021740-01). We thank Hui Pan and Jonathan Dreyfuss (Joslin Bioinformatics & Biostatistics) for analyzing bulk and single cell RNA-seq, m^6^A-seq, and eCLIP data and Manoj Gupta for mouse iPSCs. This work is supported by NIH grants R01 DK067536 (R.N.K.), UC4 DK116278 (R.N.K. and C.H.), and RM1 HG008935 (C.H). R.N.K. acknowledges support from the Margaret A. Congleton Endowed Chair and C.H. is a Howard Hughes Medical Institute Investigator. DFDJ acknowledges support by the Mary K. Iacocca Junior Postdoctoral Fellowship and American Diabetes Association grant #7-21-PDF-140.

## AUTHOR CONTRIBUTIONS

S.K. and D.F.D.J. conceived the study, designed and performed experiments, analyzed the data, and wrote the manuscript. J.W. performed mass spectrometry, and RNA and m^6^A MeRIP sequencing. N.K.B. and J.H. performed imaging and immunohistochemistry on mouse pancreatic sections. Z.Z. and J.W. performed data analysis and peak calling, and visualized the m^6^A-seq data. C.H. contributed to conceptual discussions, designed the experiments, and wrote the manuscript. R.N.K. conceived the study, designed the experiments, supervised the project, and wrote the manuscript. All the authors have reviewed, commented on, and edited the manuscript.

## STAR METHODS

### KEY RESOURCES TABLE

| REAGENT or RESOURCE | SOURCE | IDENTIFIER |
| --- | --- | --- |
| Antibodies |  |  |
| Mouse anti-METTL14<br>(1:1000) | Sigma | GN-ID4; RRID: AB_10672401 |
| Mouse anti-Ki67 (1:100) | BD Biosciences | BD550609; RRID: AB_393778 |
| Guinea Pig anti-Insulin<br>(1:400) | Abcam | ab7842; RRID: AB_306130 |
| Mouse anti-Glucagon<br>(1:10,000) | MilliporeSigma | G2654; RRID: AB_259852 |
| Rabbit anti-Somatostatin<br>(1:100) | Abcam | ab92517; RRID: AB_1143012 |
| Rabbit anti-Pdx1 (1:400) | Chemicon | AB3243 |
| Rabbit anti-β-actin (1:1,000) | Cell Signaling | 4970; RRID: AB_2223172 |
| Mouse anti-α-tubulin (1:50) | Abcam | Ab7921; RRID: AB_2241126 |
| Donkey anti-Guinea Pig<br>AF594 (1:400) | Jackson ImmunoResearch | 706-586-148; RRID:<br>AB_42340475 |
| Donkey anti-Mouse AMCA<br>(1:400) | Jackson ImmunoResearch | 715-155-150; RRID:<br>AB_2340806 |
| Donkey anti-Mouse AF488<br>(1:400) | Jackson ImmunoResearch | 715-545-150; RRID:<br>AB_2340846 |
| Donkey anti-Rabbit AF488<br>(1:400) | Jackson ImmunoResearch | 711-545-152; RRID:<br>AB_2313584 |
| Biological Samples |  |  |
| Human pancreatic sections | nPOD and the Specialized Histopathology Core at Dana-Farber/Harvard Cancer Center |  |
| Chemicals, Peptides, and Recombinant Proteins |  |  |
| DA-ZP1 | Laboratory of Amit Choudhary | n/a |
| DAPI, dilactate | Sigma | D9564 |
| TrypLE | Thermo Fisher Scientific | 12604-013 |
| FBS | Thermo Fisher Scientific | 10437028 |
| DPBS | Thermo Fisher Scientific | 14190250 |
| D-Glucose | Sigma-Aldrich | G8769 |
| FFA-BSA | Sigma-Aldrich | 3117057001 |
| DPBS | Thermo Fisher Scientific | 14190250 |
| DMEM LG | Thermo Fisher Scientific | 10-014-CV |
| DMEM no glucose | Thermo Fisher Scientific | 11966025 |
| DMSO | Sigma-Aldrich | D2650-100 |
| Penicillin-streptomycin | Corning | 30-002-CI |
| 4% PFA | Wako | 163-20145 |
| Antibody diluent | Abcam | Ab64211 |
| TRIzol reagent | Invitrogen | 15596026 |
| Critical Commercial Assays |  |  |
| MycoAlert Mycoplasma Test | Lonza | LT07-318 |
| Kit |  |  |
| ApopTag Peroxidase In Situ Apoptosis Detection Kit | MilliporeSigma | S7100: RRID: AB_2661855 |
| Qiagen RNeasy kit | Qiagen | 74104 |
| Software and Algorithms |  |  |
| Prism v.7.0 | GraphPad Software | http://www.graphpad.com;<br>RRID: SCR_002798 |
| Flowjo-v10 | FlowJo Software | http://www.flowjo.com;<br>RRID:SCR_008520 |
| ImageJ | ImageJ Software | https://imagej.net;<br>RRID:SCR_003070 |
| R version 4.1.0 | R Software | https://www.r-project.org/ |

## RESOURCE AVAILABILITY

### Lead Contact

Further information and requests for resources and reagents should be directed to and will be fulfilled by the Lead Contact, Dr. Rohit N. Kulkarni.

### Materials Availability

This study did not generate new unique reagents.

## Data and Code Availability

m^6^A-seq and RNA-seq data in differentiated hESCs have been deposited into the National Center for Biotechnology Information’s Gene Expression Omnibus under accession code no. GSE236325. RNA-seq data in METTL14 iKD embryoid bodies derived from hESCs have been deposited under accession code no. GSE236322. RNA-seq in FACS-sorted β-like cells derived from hESCs have been deposited under the accession code no. GSE236323. eCLIP data performed in human fetal EndoC-βH1 beta cells have been deposited under the accession code no. GSE236324. All data reported in this paper will be shared by the lead contact upon request. This paper does not report original code needed to reanalyze the data generated by this study. Any additional information required to reanalyze the data reported in this paper is available from the lead contact upon request.

## EXPERIMENTAL MODEL AND SUBJECT DETAILS

### Mouse studies

Ins1^Cre^, Ngn3^Cre^, and Pdx1^Cre^ Mettl14 KO mice were used for this study. Mice were maintained on a chow diet (PicoLab® mouse diet 20 – 5058). Sample sizes for animal experiments were chosen on the basis of experience in previous in-house studies of metabolic phenotypes and to balance the ability to detect significant differences with minimization of the numbers of animals used in accordance with NIH guidelines.

**Ins1^Cre^ Mettl14 KO mice:** β-cell specific Mettl14 KO mice were generated as described previously (De Jesus et al., 2019) by breeding a *Mettl14* floxed mouse with B6(Cg)-Ins1tm1.1(cre)Thor/J (Thorens et al., 2015) (Jackson Labs, USA).

**Ngn3^Cre^ Mettl14 KO mice:** Endocrine-specific Mettl14 KO mice were generated by breeding a Mettl14 floxed mouse with Tg(Neurog3-cre)C1Able (Schonhoff et al., 2004).

**Pdx1^Cre^ Mettl14 KO mice:** Endocrine-specific Mettl14 KO mice were generated by breeding a Mettl14 floxed mouse with B6.FVB-Tg(Pdx1-cre)6Tuv/J (Hingorani et al., 2003) (Jackson Labs, USA).

### Cell culture

MEL1 and H1 hESCs, miPSC, and immortalized EndoC-βH1 cell line were used to performed experiments described in this study.

### hESCs

We used MEL1 and H1, two different genetic backgrounds of hESCs. MEL1 hESCs were obtained from Murdoch Children’s Research Institute (Parkville, Victoria, Australia), and H1 (WA-01) hESCs were from WiCell (Madison, WI, USA). These hESCs were cultured on Vitronectin (VTN-N, Gibco) coated tissue culture plates in Essential 8 medium (Gibco, A1517001). Cells were split using 0.5 mM EDTA at 1:10-1:20 ratio every 4-6 days. The cells were routinely tested for mycoplasma contamination.

### miPSCs

Mouse induced pluripotent stem cells were generated as previously described (Gupta et al., 2018) and maintained in a 2i-media feeder-free system.

### EndoC-**β**H1 cells

EndoC-βH1 cell lines were obtained from Univercell-Biosolutions (France). Culture plates were coated with DMEM (glucose 4.5 g L-1; Gibco) containing fibronectin (2 μg mL-1; Gibco), and extracellular matrix (1% vol vol-1; Sigma) for at least an hour in 5% CO2 at 37°C. EndoC-βH1 cells were grown on coated 6-well plates containing DMEM (glucose 1 g L-1), BSA fraction V (2% wt vol-1) (Roche), 2-mercaptoethanol (50 μM; Sigma), nicotinamide (10 mM; Sigma), transferrin (5.5 μg mL-1; Sigma), and sodium selenite (6.7 ng mL-1; Sigma).

## METHODS DETAILS

### Immunohistochemistry

Mouse pancreas was collected and fixed in 4% formaldehyde at 4 °C overnight, followed by paraffin embedding. Human pancreas sections were obtained from the nPOD and the Specialized Histopathology Core at Dana-Farber/Harvard Cancer Center. Five-micron-thick slides were cut and subjected to immunostaining. Slides were heated in Tris-EDTA buffer (10mM Tris base, 1 mM EDTA, 0.05% Tween 20, pH 9.0) followed by blocking with donkey serum and incubated with primary antibodies against METTL14 (sigma HPA038002, 1:1000) and insulin (Abcam, ab7842, 1:400). Slides were heated in 10 mM sodium citrate, followed by blocking with donkey serum and incubated with primary antibodies against insulin (Abcam, ab7842, 1:400), glucagon (MilliporeSigma, G2654, 1:10,000), Somatostatin (ab64053, 1:100), Ki67 (BD550609, 1:100), Pdx1 (Chemicon, AB3243, 1:400), and TUNEL (ApopTag, Chemicon, S7100) and counterstained with DAPI (MilliporeSigma, D9564, 1:6,600). Images were captured using a Zeiss Axio Imager A2 upright fluorescence microscope. METTL14 mean fluorescence intensity (MFI) was measured in insulin positive areas using ImageJ 1.51s. The β-cell mass was calculated by multiplying the pancreas weight of the mouse with the ratio of the insulin-positive area to the pancreatic tissue area. For estimation of β-cell proliferation, ∼1,500 cell nuclei on average were counted per section, and data were expressed as percentage of Ki67+Ins+ cells. To assess cell death, apoptotic index was measured by quantification of the percentage of TUNEL+Ins+ cells.

### Protein isolation and western blotting

Total protein amounts were collected from tissue and cell line lysates using RIPA buffer with proteinase and phosphatase inhibitors (Sigma). Protein concentrations were determined using the BCA method followed by standard western immunoblotting of proteins using different primary antibodies: Anti-METTL14 (HPA038002, Sigma), Anti-β-actin (4970, Cell Signaling), and Anti-α-tubulin (7291, abcam). The blots were developed using chemiluminescent substrate ECL and quantified using ImageJ 1.15s.

### In vitro pancreatic differentiation

hESC colonies were dissociated into single cells using TrypLE (Gibco, 12604-021) and reseeded in VTN-N coated plates in E8 medium containing 5 μM Y-27632 (Fisher, NC1286855). Differentiation was initiated 24 h to 48 h after plating when the culture was 90% in confluency.

Cultures were rinsed with DPBS without Mg2+ and Ca2+ (Gibco) and differentiation medium was added and refreshed every day as reported previously (Kahraman et al., 2022). MCDB 131 medium with 10 mM glucose and 1% Glutamax was supplemented with: on day 1– 0.5% FFA-BSA, 1.5 g/L NaHCO_3_, 100 ng/ml GDF8, 3 μM CHIR-99021; on day 2 – 0.5% FFA-BSA, 1.5 g/L NaHCO_3_, 100 ng/ml GDF8, 0.3 μM CHIR-99021; on day 3 – 0.5% FFA-BSA, 1.5 g/L NaHCO_3_, 100 ng/ml GDF8; on days 4-6 – 2% FFA-BSA, 1.5 g/L NaHCO_3_, 0.25 mM Ascorbic acid, 1:50000 ITS-X, 50 ng/ml FGF7; on days 7-8 – 2% FFA-BSA, 2.5 g/L NaHCO_3_, 0.25 mM Ascorbic acid, 1:200 ITS-X, 50 ng/ml FGF7, 200 nM TPB, 0.25 μM SANT-1, 1 μM Retinoic Acid, 100 nM LDN-193189; on days 9-13 – 2% FFA-BSA, 2.5 g/L NaHCO_3_, 0.25 mM Ascorbic acid, 1:200 ITS-X, 50 ng/ml FGF7, 0.25 μM SANT-1, 0.1 μM Retinoic Acid; on days 14-16 – 20 mM final glucose, 2% FFA-BSA, 2 g/L NaHCO_3_, 1:200 ITS-X, 0.25 μM SANT-1, 0.05 μM Retinoic Acid, 100 nM LDN-193189, 1 μM T3, 10 μM ALK5 inhibitor II, 10 μM ZnSO4, 10 μg/ml Heparin; on days 17-23 – 20 mM final glucose, 2% FFA-BSA, 2 g/L NaHCO_3_, 1:200 ITS-X, 100 nM LDN-193189, 1 μM T3, 10 μM ALK5 inhibitor II, 10 μM ZnSO_4_, 100 nM GSiXX; and on days 24-30 – 20 mM final glucose, 2% FFA-BSA, 2 g/L NaHCO_3_, 1:200 ITS-X, 1 μM T3, 10 μM ALK5 inhibitor II, 10 μM ZnSO_4_, 1 mM N-Cys, 10 μM Trolox, 2 μM R428. Mouse iPSCs were differentiated into pancreatic β-like cells as described previously (Gupta et al., 2018; Liu and Lee, 2012).

Pancreatic β-like cells were harvested on day 8 for total RNA isolation and transcript analyses of β-cell developmental markers.

### Liquid chromatography–mass spectrometry quantification of m^6^A

Total RNA was isolated by TRIzol reagent and mRNAs were purified by Dynabeads mRNA purification kit two times, followed by rRNA depletion with RiboMinus™ Eukaryote Kit v2kit. The purified mRNAs were digested with nuclease P1 (Sigma, N8630) for 2 h at 42°C, and then with FastAP Thermosensitive Alkaline Phosphatase (Thermofisher Scientific, EF0651) for 4 h at 37°C. The samples were then filtered (0.22 mm, Millipore) and injected into a C18 reverse phase column coupled online to an Agilent 6460 LC–MS/MS spectrometer. The nucleosides were quantified using retention time and the nucleoside to base ion mass transitions (268-to-136 for A; 282-to-150 for m^6^A). Quantification was performed by comparing this with the standard curve obtained from nucleoside standards run with the same batch of samples.

### m^6^A immunoprecipitation and sequencing

Total RNA was isolated by TRIzol reagent and mRNAs were purified by Dynabeads mRNA purification kit. Purified mRNA was fragmented by Bioruptor® Pico Sonication System and input was saved before m^6^A immunoprecipitation. m^6^A immunoprecipitation was performed with EpiMark®N6-Methyladenosine Enrichment Kit (NEB, E1610S) following the manufacturer protocol. Then, RNA libraries were prepared for both input and IP samples using TruSeq® Stranded mRNA Library Prep (Illumina, 20020594) following the manufacturer protocol.

Sequencing was performed at the University of Chicago Genomics Facility on an Illumina NovaSeq 6000 machine.

### Differential methylation analysis for m^6^A-seq

We used the MeRIP R objects which contain mapped read counts in 50-bp bins of each gene. We then performed peak calling, peak merging, and read counting in merged peaks using the R package MeRIPtools (Zhang et al., 2020). The starting MeRIP objects for this analysis use the Ensembl gene annotation (version 94). We performed m^6^A profiling analysis of count data using the R package DESeq2, which is one of the methods used by MeRIPtools and fits the count data to a negative binomial model (Love et al., 2014). We first filtered out the peaks that had a total read count less than 10 across all samples. Using Wald tests, we then tested for significant differences in m^6^A enrichment between developmental stages.

### Differential expression analysis

We performed RNA-seq analysis of count data using the R package DESeq2, which is one of the methods used by MeRIPtools and fits the count data to a negative binomial model (Love et al., 2014). We first filtered out the genes that had a total read count less than 10 across all samples. Using Wald tests, we then tested for significant differences in gene expression between developmental stages.

### Generation of inducible METTL14 knockdown hESC lines

H1 and MEL1 hESCs cells were transduced using high titer lentiviral particles in the presence of 10 μg/ml polybrene. SMARTvector human lentiviral vectors containing shRNAs targeting METTL14 (iKD1; V3SH7669-224822773, iKD2; V3SH7669-225341368, iKD3; V3SH7669-229883026) were purchased from Dharmacon. Non-targeting control lentiviral particles were used as scramble control (iSCR; VSC10712). Puromycin (2 μg/ml) was added to the culture media starting from post-transduction day 4 to select stable inducible knockdown hESCs.

Doxycycline was added to culture medium at 2 μg/ml concentration to induce shRNA expression.

### Mettl14 knockdown in miPSCs

Briefly, miPSCs maintained on a 2i system were differentiated into pancreatic β-like cells as described previously (Gupta et al., 2018; Liu and Lee, 2012). At time 0h of differentiation, cells were mixed with Lipofectamine RNAiMAX Reagent (Life Technologies) and small interfering RNA complexes (Dharmacon) at a final concentration of 15 nmol/L siRNA according to manufacturer instructions. Media was exchanged 6h post-transfection and differentiated β-like-cells were collected at day 8 of differentiation. Dharmacon siGENOME Non-Targeting siRNA Control Pools (D-001206-13-05) and siGENOME Mouse Mettl14 siRNA (M-063715-00-0005) were used.

### EB formation assay using miPSCs

Embryonic bodies were generated as previously described (Gupta et al., 2018). Briefly, miPSCs grown in a 2i system were collected using Accutase (Invitrogen), and two million iPSCs were seeded in 10 cm petri-dishes containing high glucose DMEM supplemented with 20% FBS. Media was replaced every 24h, and cells started to form EBs at day 2 of differentiation. EBs were then harvested, washed with DPBS, and lysed in TRIzol for RNA isolation using RNeasy Kit (Qiagen) according to the manufacturer instructions.

### EB formation assay using hESCs

H1 and MEL1 hESCs were dissociated into single cells and 500 cells per microwell were seeded in AggreWell™400 24 well plates according to the manufacturer instructions. AggreWell™ EB Formation Medium – supplemented with doxycycline (2 μg/ml) to induce shRNA expression – was refreshed every other day for ten days. EBs were then harvested, washed with DPBS, and lysed in TRIzol for RNA isolation using RNeasy Kit (Qiagen) according to the manufacturer instructions.

### FAC-Sorting insulin-expressing **β**-like cells

Cells were harvested using TrypLE and neutralized in DMEM containing 10% FBS. The cell pellet was resuspended in DA-ZP1 containing cell media and incubated in 37°C for 30 min. Cells were washed with DPBS and the cell pellet was resuspended in fresh DA-ZP1–free media. Cells were sorted by Aria as described previously (Kahraman et al., 2021). Sorted cells were washed with DPBS and lysed in TRIzol for RNA isolation using RNeasy Micro Kit (Qiagen) according to the manufacturer instructions.

### RNA-seq and data analysis

RNA libraries from METTL14 iKD EB were prepared using TruSeq Stranded Total Library Prep (Illumina) and RNA libraries from METTL14 iKD β-like cells were prepared using SMARTer Stranded Total RNA-Seq kit v2 (Takara) following the manufacturer protocol. Sequencing was performed on an Illumina NovaSeq 6000 according to the manufacturer instructions.

Approximately 50 million paired-end 100-bp reads were generated for each sample. We aligned the adapter-trimmed reads to the human transcriptome using Kallisto, converted transcript counts to gene counts using tximport, normalized the counts by trimmed mean of M-values (TMM) (Robinson and Oshlack, 2010), and transformed normalized counts into log2 counts per million (logCPM) with Voom (Law et al., 2014). Pathway analysis was done using the ConsensusPathDB interaction database (Herwig et al., 2016).

### Single cell RNA-seq analysis of GSE114412

We downloaded this previously published dataset (Veres et al., 2019) from the Gene Expression Omnibus (GEO). We removed genes that have average counts of 0. Similar cells were clustered together using a graph-based clustering algorithm and the data was then normalized.

### Enhanced crosslinking and immunoprecipitation (eCLIP) assay

The eCLIP assays were performed by Eclipse Bioinnovations using UV-crosslinked EndoC-βH1 cells following the protocol detailed in (Van Nostrand et al., 2016). For these assays, the anti-METTL14 antibody A305-847A (Bethyl Laboratories) was used, and two independent biological replicates were performed. All comparisons were done relative to the size-matched input control. Sequences were processed and mapped using the pipeline described in (Van Nostrand et al., 2016). We bin the aligned reads into 200-bp bins for all IP and input samples. For each IP and its matched input sample, we retained only those bins with one or more reads mapping. We detected genome regions of RNA enrichment using Piranha (Uren et al., 2012) by zero-truncated negative binomial regression, with each IP sample as response and the matched input sample as covariate (i.e. the baseline). We imported peaks data and only kept peaks that had p-values of at least 0.05. We annotated the overlapped peaks that were within 5000-bp up/down-stream of the nearest genes.

## QUANTIFICATION AND STATISTICAL ANALYSIS

Statistical analyses were performed using GraphPad Prism software version 7.0a (GraphPad Software Inc., La Jolla, CA). The results are expressed as the mean ± standard error of the mean. Specific statistical tests for each experiment are described in the figure legends. In figures * p < 0.05, **p < 0.01 and ***p < 0.001. Data represent mean ± SEM; *p < 0.05; **p < 0.01; ***p < 0.001.

---

**Figure S1, related to Figure 2: Changes in gene expression profiles during in vitro β-cell differentiation.**

**(A)** Gene markers of posterior foregut (S3). Multiple t-test vs S0 (H1 n = 2 and MEL1 n = 2 independent biological replicates).

**(B)** Gene markers of pancreatic progenitors (S4). Multiple t-test vs S0 (H1 n = 2 and MEL1 n = 2 independent biological replicates).

**(C)** Gene markers of cells at S6. Multiple t-test vs S0 (H1 n = 2 and MEL1 n = 2 independent biological replicates).

**(D)** m^6^A enriched peaks (H1, S0, S3, S4, S6, n = 2 biological replicates).

**(E)** m^6^A enriched peaks (MEL1, S0, S3, S4, S6, n = 2 biological replicates).

**(F)** m^6^A distribution in S0, S3, S4, S6 samples (H1 n = 2 and MEL1 n = 2 biological replicates).

**Figure S2, related to Figure 3: eCLIP assay in EndoC-βH1.**

**(A)** Interaction of GO terms and genes enriched in eCLIP analysis in EndoC-βH1 (https://toppcluster.cchmc.org/).

**(B)** Coverage plot for PDX1.

**(C)** Coverage plot for MNX1.

**(D)** Coverage plot for FOXA2.

**(E)** Coverage plot for HNF1A.

**(F)** Coverage plot for PCNT.

**Figure S3, related to Figure 3: In vitro differentiation of miPSCs into β-like cells.**

**(A)** In vitro differentiation protocol for mouse iPSCs towards β-like cells.

**(B)** RT-PCR analysis of β-like cells derived from miPSCs (β-like cells n = 4; miPSCs n = 3; mouse islets n = 3 independent biological samples). Expression was normalized against the mean level in undifferentiated miPSCs.

