## Supplementary figures and images for "m^6^A mRNA Methylation Regulates Early Pancreatic β-Cell Differentiation"

### Supplemental Figures

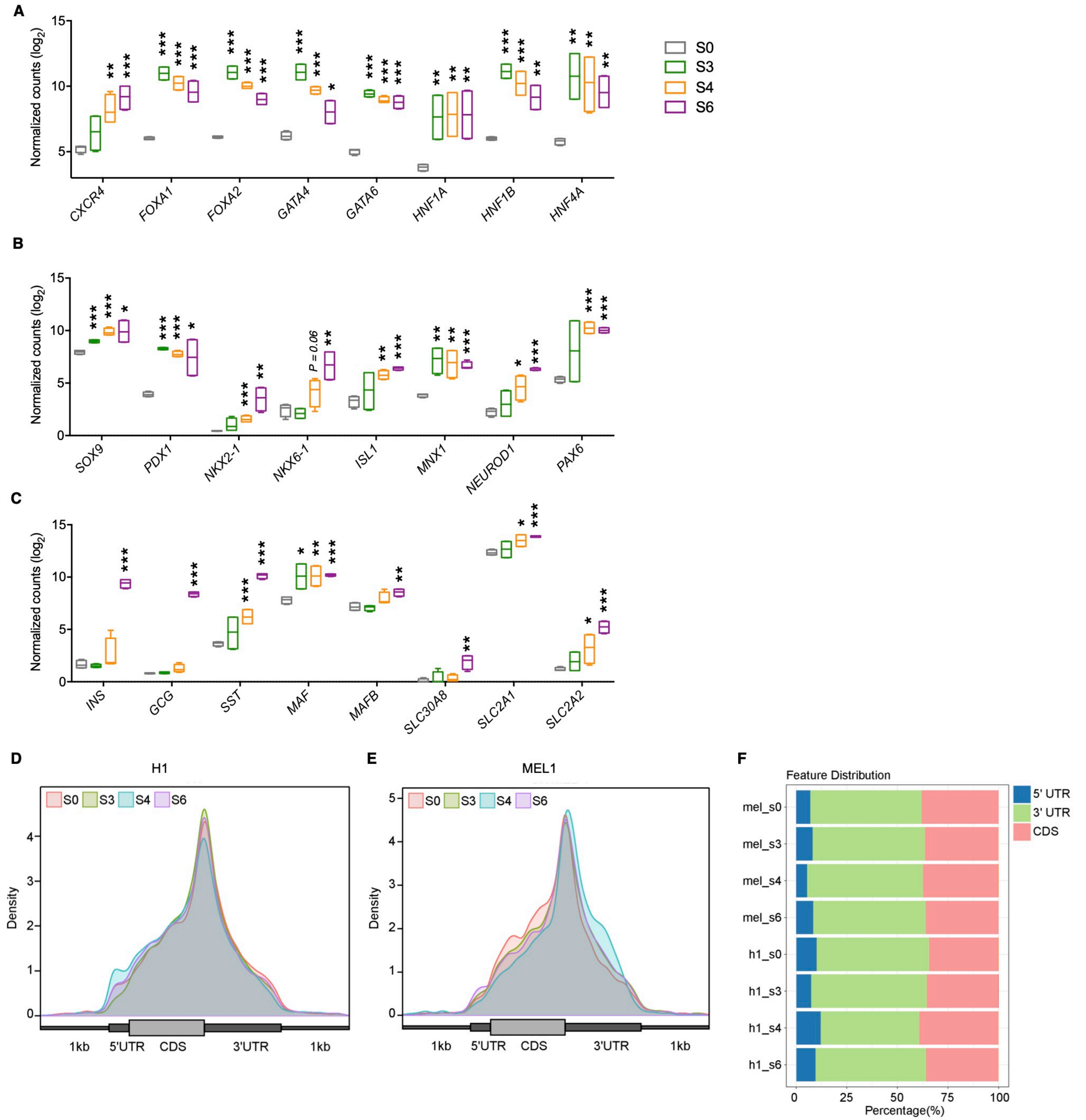

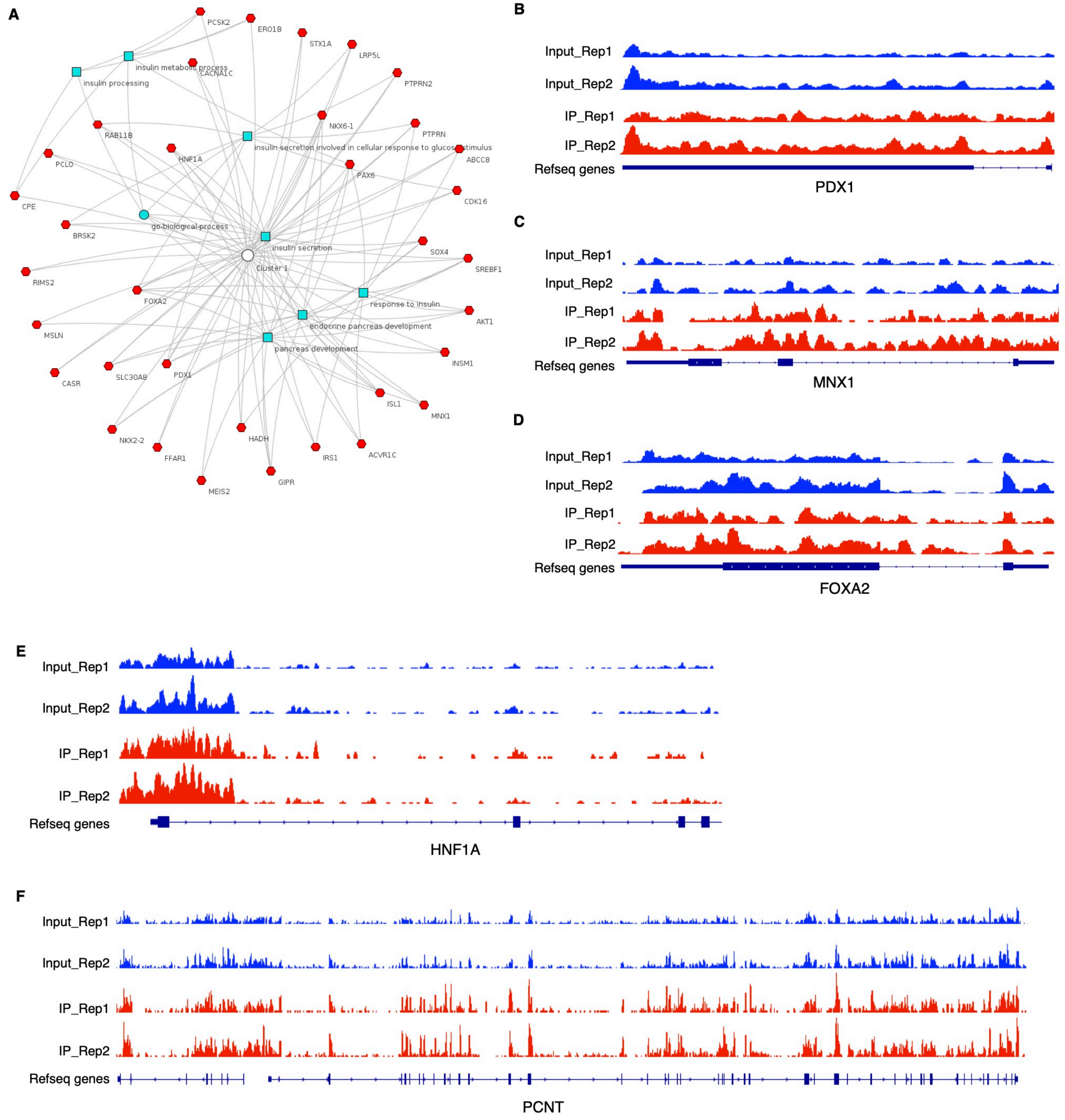

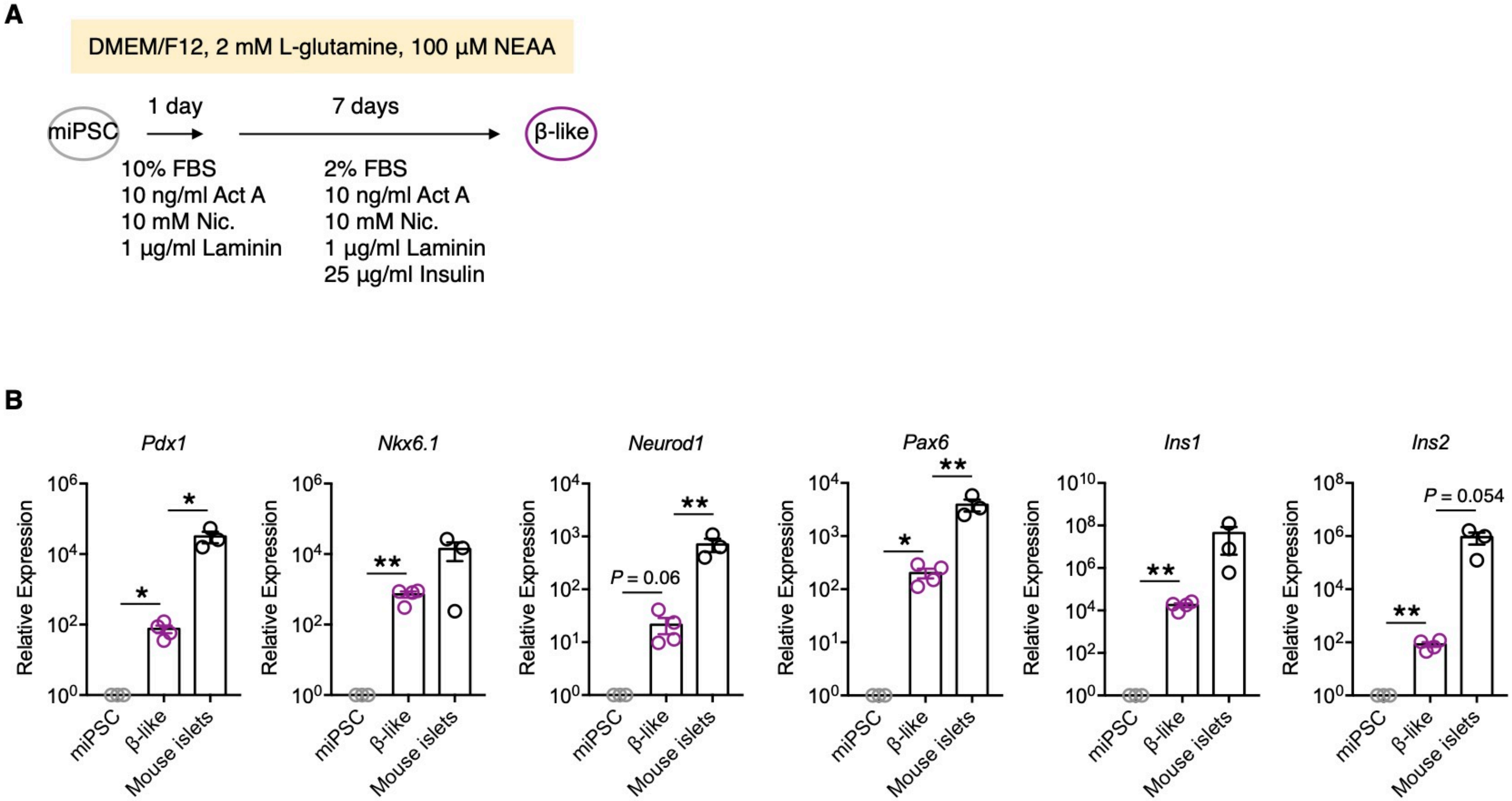
